# Immunoregulatory gene *GIMAP6* suppresses lethal atherosclerotic vasculopathy and ischemic heart failure

**DOI:** 10.64898/2026.01.26.701509

**Authors:** Chen Xiang, Kaja Arusha, Danielle Springer, Sachiko Nakamori, Ping Du Jiang, Zhi-Hong Yang, Jing Cui, Yu Zhang, Huie Jing, Ann Y. Park, Maria H. Zhu, Sarah E. Weber, Deniz Cagdas, Hassan Abolhassan, Nasrin Behniafard, Margery G. Smelkinson, Qihui Liang, Elif Everest, Julia F. Kun, Alyssa Grogan, Jennifer D. Treat, Christa S. Zerbe, Steven M. Holland, Renu Virmani, Alan T. Remaley, Helen C. Su, Lixin Zheng, Michael J. Lenardo

## Abstract

Controlling hyperlipidemia has reduced but not eliminated atherosclerotic cardiovascular disease as a predominant cause of human mortality. Here, we report that loss of the immunoregulatory gene GTPase of immunity-associated protein 6 (*GIMAP6*), causes an inflammatory vasculopathy and accelerated atherosclerosis in the absence of hyperlipidemia. These pathologic changes in turn result in progressive cardiac ischemia, myocardial infarction, and heart failure, culminating in early death. In humans, rare deleterious GIMAP6 variants are associated with premature severe cardiovascular disease. These findings reveal GIMAP6 to play an important protective role against atherosclerotic cardiovascular disease whose identification offers opportunities for improved risk management and a target for new therapies.

## INTRODUCTION

Despite the effectiveness of cholesterol- and low-density lipoprotein (LDL)-lowering therapies, about half of clinically relevant atherosclerotic cardiovascular disease (ASCVD) events occur in people without severe hyperlipidemia, pointing to the existence of additional significant risk factors.^1-6^ The landmark CANTOS (Canakinumab Anti-Inflammatory Thrombosis Outcomes Study) trial provided evidence that inflammation is one causal driver of this added risk, as anti-inflammatory therapy targeting interleukin-1β reduced cardiovascular events independent of lipid levels.^7^ Formation of macrophage foam cells, which is pathognomonic for ASCVD, is now understood to reflect not only abnormal lipid deposition but also a key amplifying event, as these specialized macrophages actively secrete cytokines, recruit other immune cells, and propagate the chronic vascular inflammation that drives plaque progression.^6^ Whether molecular changes that increase the lipid uptake in macrophages without the drive of excessive plasma LDL and cholesterol could lead to the same consequence of foam cell formation, ASCVD, and ischemic heart failure is currently unclear.

GTPase of immunity-associated protein 6 (GIMAP6) is a member of a linked family of GIMAP genes conserved in vertebrates, but its functions remain largely undefined.^8^ Deficiency of GIMAP6 in humans is now recognized as an Inborn errors of immunity (IEI), causing a complex immunodysregulation that includes altered T cell subsets, abnormal immunoglobulin levels with autoantibodies, lymphadenopathy, splenomegaly, recurrent infections, vasculitis and subcortical ischemic lesions.^9^ Similarly, studies in *Gimap6* knockout mice reveal that the protein has important roles in autophagy, T cell homeostasis, and antibacterial immunity.^9^ Unexpectedly, these *Gimap6*-deficient mice also exhibit premature death under specific-pathogen-free conditions without evidence of infection, a lethal phenotype previously attributed to a microangiopathic kidney abnormality of unknown cause.^9^ Several lines of evidence now point to an underlying vasculopathy as the cause of death. GIMAP6 and the related protein GIMAP5 help control cellular lipid homeostasis,^9,10^ and notably, genome-wide association studies (GWAS) show a high-confidence association between the *GIMAP6* locus and C-reactive protein, a key clinical biomarker for cardiovascular inflammation.^11,12^ Furthermore, the link between the GIMAP family and vascular inflammation is supported by its identification as a susceptibility locus for Behcet’s disease, a systemic vasculitis.^13^ Given the clinical observation of vasculopathy in humans, the unexplained premature death in mice, and the protein’s connection to lipid metabolism and inflammatory markers, we further investigated the pathophysiology of GIMAP6 deficiency.

In this study, we characterize the lethal inflammatory vasculopathy in *Gimap6*^−*/*−^ mice, demonstrating that GIMAP6 deficiency leads to severe ASCVD that does not require but may be exacerbated by hyperlipidemia and culminates in fatal myocardial infarction, heart failure, and early death. This genetic effect is accelerated in females. GIMAP6 acts within macrophages to limit pathological lipid accumulation and foam cell formation. We also observing various vascular abnormalities, including stroke, in individuals with rare GIMAP6 deficiencies. Thus, our work identifies an immunodeficiency gene that controls atherosclerosis by a new mechanism that underlies inflammation-driven vasculopathy and ASCVD.

## RESULTS

### *Gimap6* deficiency in mice drives lethal spontaneous atherosclerosis and heart failure independent of hyperlipidemia

To explore our preliminary observation of premature death in *Gimap6*^−*/*−^ mice,^9^ we conducted a long-term survival study. We found that the lethal phenotype is fully penetrant on the inbred C57BL/6 background, as all *Gimap6*^−*/*−^ mice died prematurely (Figure 1A) with a median survival of 31 weeks for males and only 21 weeks for females (Figure 1B). Heterozygous female *Gimap6*^*+/*−^ mice, a cohort that included breeders, showed only a modest reduction in survival (Figure 1A). Euthanasia of moribund mice revealed striking cardiomegaly that worsened in the weeks preceding death (Figures 1C).

**Figure 1.**
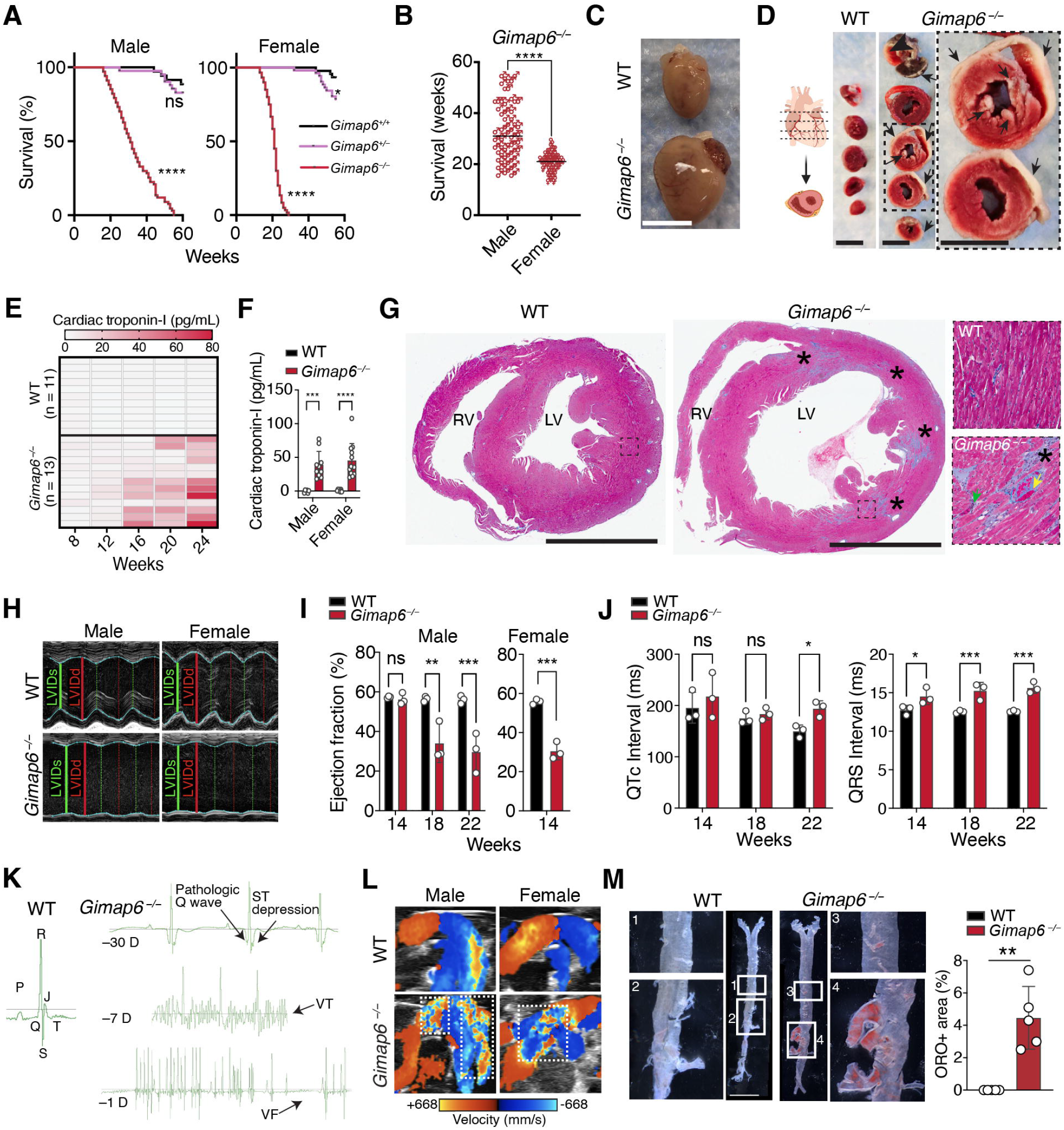
*Gimap6* deficiency leads to premature death due to myocardial infarction and atherosclerosis. (A) Kaplan-Meier survival curves for *Gimap6*^−*/*−^ (males, n = 67; females, n = 60), *Gimap6*^*+/*−^ (males, n = 41; females, n = 53), and *Gimap6*^*+/+*^ (males, n = 35; females, n = 47) mice on a standard-fat diet (SFD). (B) Age at death (weeks) for male (n = 67) and female (n = 60) Gimap*6*^−*/*−^ mice shown in (A). Each point represents an individual animal. (C) Representative images of gross heart anatomy from 5-month-old male wild-type (WT) and *Gimap6*^−*/*−^ mice. Scale bar, 0.5 cm. (D) Left: diagram of transverse heart sections. Right: 2,3,5-Triphenyltetrazolium chloride-stained sections from 7-month-old male WT and *Gimap6*^−*/*−^ mice appearing moribund (severely reduced activity, hunched posture, ruffled fur). Pale non-staining areas (black arrows) indicate infarcted tissue, viable myocardium stains red. The dashed black box highlights a 2.5x enlarged view of selected sections. Scale bar, 0.5 cm. (E) Heatmap displaying plasma cardiac troponin I (pg/mL) levels in individual WT (n = 11) and *Gimap6*^−*/*−^ (n = 13) SFD-fed mice measured from blood samples collected every 4 weeks between 8 and 24 weeks of age (n = 3–7 per time point, pooled from 3 independent experiments). Each row represents a single mouse. (F) Quantitation of plasma cardiac troponin I in male (n = 10) and female (n = 12) *Gimap6*^−*/*−^ mice appearing moribund compared to age- and sex-matched healthy WT controls (male, n = 5; female n = 6). Each dot represents an individual mouse. Bars indicate mean ± standard deviation (SD). (G) Representative Masson’s trichrome-stained transverse heart sections from 5-month-old male WT and *Gimap6*^−*/*−^ mice. The right ventricle (RV) and left ventricle (LV) are labeled. Healthy myocardium appears pink and asterisks (*) highlight fibrotic tissue (blue) indicating areas of potential prior infarcts. The magnified inset (6x) from the *Gimap6*^−*/*−^ heart further highlights the presence of inflammatory cell infiltrates (green arrow) and vascular abnormalities (yellow arrow) within the fibrotic regions. Scale bar, 0.25 cm. (H) Representative short-axis 2D M-mode echocardiograms from male WT and *Gimap6*^−*/*−^ mice at 22 weeks and female WT and *Gimap6*^−*/*−^ mice at 14 weeks. The overlaid curve (blue) traces the left ventricular internal diameter (LVID) with bars indicating systole (LVIDs, green) and diastole (LVIDd, red). (I) Ejection fraction (EF) values for WT and *Gimap6*^−*/*−^ mice computed from echocardiograms (n = 3 per group, per time point). (J) The corrected QT interval (QTc), and QRS wave duration measurements from male WT (n = 3) and *Gimap6*^−*/*−^ (n = 3) mouse hearts at 14, 18, and 22 weeks. Data are presented as mean ± SEM, with each dot representing an individual mouse. (K) Representative telemetry electrocardiogram (EKG) recordings from one female WT and one female *Gimap6*^−*/*−^ mouse. The WT tracing shows normal P-QRSJ-T waves. *Gimap6*^−*/*−^ recordings display progressive abnormalities over the 30-day observation period. Ventricular fibrillation (VF), ventricular tachycardia (VT). (L) Color Doppler echocardiogram of the aortic arch during diastole in 3.5-month-old WT and *Gimap6*^−*/*−^ male and female mice. Colors represent blood flow velocity: red indicates flow toward the transducer, blue indicates flow away from the transducer. The color scale represents velocity from −668 mm/s (blue) to +668 mm/s (red). Dotted boxes highlight regions of turbulent flow. (M) Representative en face images of Oil Red O (ORO)-stained aortae from cohoused 5-month-old male WT and *Gimap6*^−*/*−^ mice maintained on an SFD. Insets (1–4) show 3.5x magnified views of the indicated regions. Scale bar, 0.5 cm. Right: quantification of ORO-positive plaque area, expressed as a percentage of the total aortic surface area. Each point represents an individual animal. *P* values were calculated by log-rank (Mantel-Cox) test in (A), multiple unpaired *t*-tests, with the Holm-Šídák method used to correct for multiple comparisons in (I) and (J), or unpaired *t*-test in (B) (F) and (M). Statistical significance is denoted as not significant (ns), \**P* < 0.05, \*\**P* < 0.01, \*\*\**P* < 0.001, \*\*\*\**P* < 0.0001.

The combination of progressive cardiac hypertrophy and sudden decompensation led us to hypothesize that underlying cardiovascular disease caused the premature mortality in *Gimap6*^−*/*−^ mice. We therefore carried out triphenyl tetrazolium chloride staining of cardiac tissues sections from moribund animals that revealed extensive areas of infarction in *Gimap6*^−*/*−^ hearts, as marked by pale unstained regions characteristic of tissue death (Figure 1D).^14^ Also, plasma cardiac troponin I levels began rising as early as 8 to 12 weeks of age in deficient animals (Figure 1E), consistent with subclinical injury, and a marked elevation in all moribund mice indicated terminal myocardial damage (Figure 1F). Histopathological examination with Masson’s trichrome staining revealed prominent ventricular dilation, vascular abnormalities, and extensive myocardial fibrosis in *Gimap6*^−*/*−^ mice, with large fibrotic areas replacing healthy myocardium (Figure 1G). High-magnification views confirmed this severe pathology, showing that in contrast to the organized myocardium of wild-type controls, the tissue in knockout mice was extensively replaced by fibrotic collagen and infiltrated by inflammatory cells, leading to disorganized cardiomyocytes (Figure 1G, enlarged panel). Together, the results suggest that cardiac damage in *Gimap6*^−*/*−^ mice, along with compensatory inflammation and tissue remodeling,^15^ was ongoing prior to heart failure and death.

In accord with the pathological data, M-mode echocardiographic assessments demonstrated progressive left ventricular dilatation and rapidly declining cardiac contractility over time in male and female *Gimap6*^−*/*−^ mice (Figure 1H). The ejection fraction declined significantly in males from 14 to 22 weeks of age and was markedly reduced in females as early as 14 weeks, indicating accelerated and severe heart failure in both sexes, with a markedly earlier and more severe onset in females (Figure 1I). Moreover, Doppler echocardiography of the aortic arch in *Gimap6*^−*/*−^ mice revealed turbulence and distortion of aortic laminar flow, suggestive of vascular damage or early atherosclerotic changes (Figure 1L).^16^ To determine if the fatal cardiac phenotype was driven by underlying vascular disease, we performed en face Oil Red O (ORO) staining to visualize atherosclerotic plaques in the aorta. Standard C57BL/6 mice are known to be resistant to atherosclerosis and do not develop spontaneous plaques on a standard-fat diet (SFD).^17^ Remarkably, even on an SFD, *Gimap6*^−*/*−^ mice developed significant spontaneous atherosclerotic plaque deposition, a feature absent in WT littermates. Quantitative analysis showed these plaques covered about 4% of the aortic surface in knockout mice. The lesions were most severe at vessel branch points especially at the junction of the renal arteries consistent with classic atherosclerotic plaques (Figure 1M).^18^ The presence of atherosclerotic plaques can explain not only the previously observed kidney damage but also the cerebrovascular ischemic events observed in a GIMAP6-deficient human patient.^9^ To determine whether the severe pathology observed in *Gimap6*^−*/*−^ mice was driven by alterations in plasma lipids, we conducted lipoprotein fractionation analysis. On an SFD, *Gimap6*^−*/*−^ mice maintained a normal, high-density lipoprotein (HDL)-dominant lipid profile, with non-significant changes in total cholesterol but a statistically significant, albeit modest, increase in triglycerides in males compared to wild-type controls (not shown). The vessel pathology was exacerbated when mice were challenged with a high-fat diet (HFD). Together, these data demonstrate that *Gimap6* deficiency alone is sufficient to cause a novel, monogenic form of spontaneous ASCVD that leads to fatal heart disease and does not require hyperlipidemia.

### *Gimap6* deficiency and hyperlipidemia synergize to exacerbate lethal ASCVD in *Ldlr*^−/−^ mice

While traditional atherosclerosis models such as *Ldlr*^−*/*−^ or *Apoe*^−*/*−^ mice require an HFD to develop extensive atherosclerosis in the aorta and large vessels, they rarely exhibit coronary artery involvement, spontaneous infarction, heart failure, or death.^19^ As expected, homozygous *Gimap6*^−*/*−^*Ldlr*^−*/*−^ mice had markedly reduced survival compared to *Gimap6*^−*/*−^ or *Ldlr*^−*/*−^ mice, while heterozygous *Gimap6*^*+/*−^*Ldlr*^−*/*−^ mice showed modestly worsened survival (Figure 2A). Moreover, female *Gimap6*^−*/*−^*Ldlr*^−*/*−^ mice had a shorter lifespan than their male counterparts (Figure 2A). Consistent with these findings, both SFD-fed and HFD-fed *Gimap6*^−*/*−^*Ldlr*^−*/*−^ mice developed markedly more severe atherosclerosis with aneurysm formation compared to *Ldlr*^−*/*−^ mice (Figures 2B-C). Disease became apparent within 2 months in females and 3 months in males, and an HFD-accelerated disease progression caused severe intramural hemorrhage and aneurysms (Figures 2B-C). Masson’s trichrome staining of *Gimap6*^−*/*−^*Ldlr*^−*/*−^ thoracic aortas revealed large aggregates of infiltrating macrophages with a lipid-laden foam cell morphology (Figures 2D, top). Abdominal aortas of *Gimap6*^−*/*−^*Ldlr*^−*/*−^ mice contained severe plaques consisting of foam cells (red arrows), inflammatory cells, and necrotic cores (black arrows) in the tunica media far worse than in matched *Ldlr*^−*/*−^ controls, which showed minimal infiltration (Figures 2D, bottom). Immunofluorescence staining confirmed that *Gimap6*^−*/*−^*Ldlr*^−*/*−^ lesions were characterized by a massive, transmural infiltration of CD68^+^ macrophages extending from the intima deep into the tunica media (Figure 2E). This was accompanied by severe destruction of the elastic lamina and surrounding smooth muscle cell layer, consistent with advanced, unstable plaques in human disease, which are not typically seen in other murine models of ASCVD (Figure 2E).^20,21^

**Figure 2.**
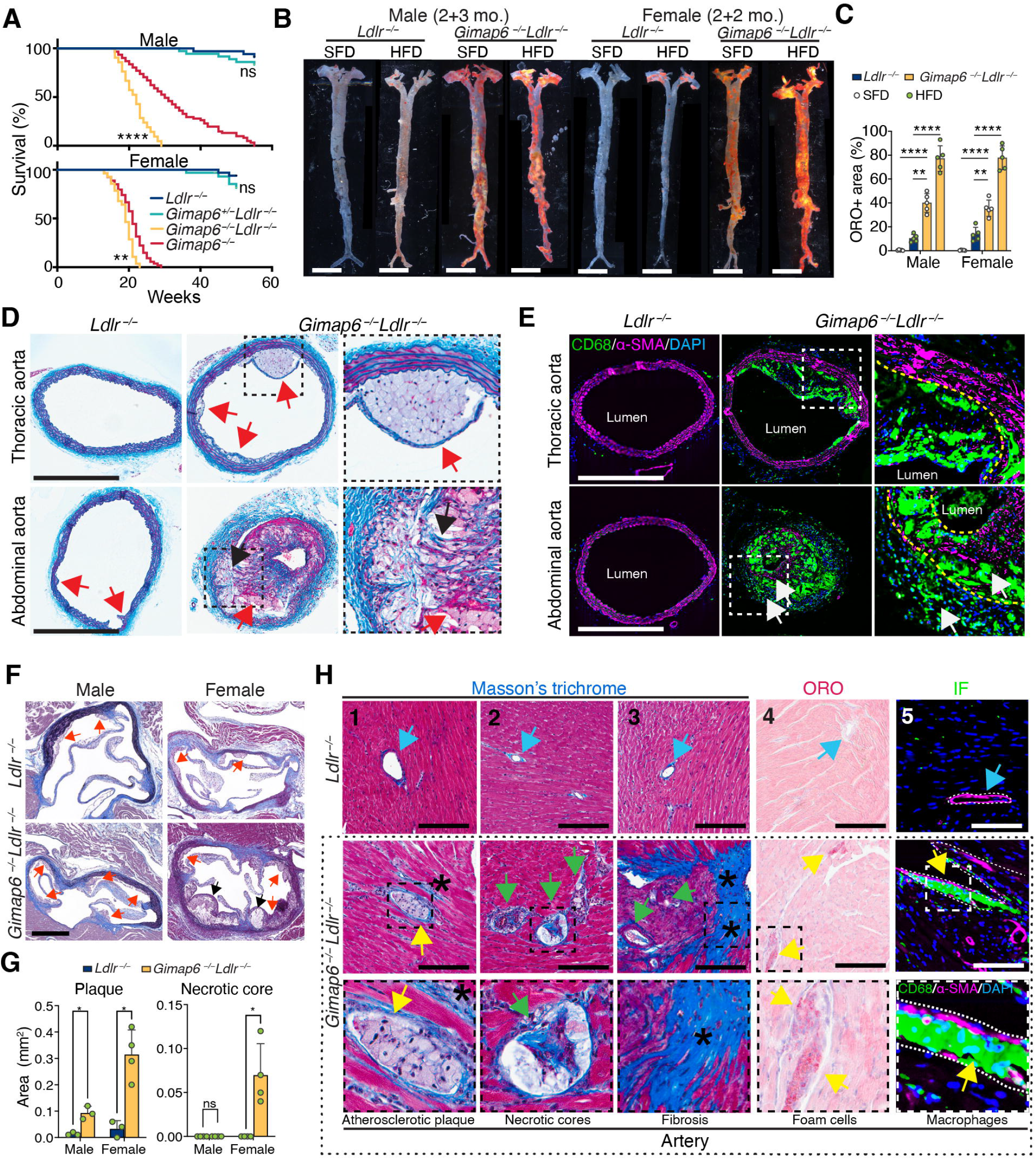
*Gimap6* deficiency synergizes with hyperlipidemia to drives lethal coronary artery disease in *Ldlr*-deficient mice. (A) Kaplan-Meier survival curves for *Ldlr*^−*/*−^ (males, n = 34; females, n = 33; teal), *Gimap6*^*+/*−^ *Ldlr*^−*/*−^ (males, n = 36; females, n = 33, blue), *Gimap6*^−*/*−^ *Ldlr*^−*/*−^ (males, n = 21; females, n = 26, yellow), *Gimap6*^−*/*−^ (males, n = 67; females, n = 60, red), mice fed a SFD. *Gimap6*^−*/*−^ control data are replotted from Figure 1A. (B) Representative en face images of Oil Red O (ORO)-stained aortae from cohoused male and female *Ldlr*^−*/*−^ and *Gimap6*^−*/*−^ *Ldlr*^−*/*−^ mice. All mice were fed an SFD for the first 2 months. Both genotypes were then assigned to either a continued SFD or switched to an HFD. Males and females were maintained for an additional 3 or 2 months, respectively, before analysis. Scale bar, 0.5 cm. (C) Quantification of ORO-positive plaque area in (B), expressed as a percentage of the total aortic surface area. Each point represents an individual animal, bars indicate mean ± SD. (D) Representative thoracic (top row) and abdominal (bottom row) aortic sections stained with Masson’s trichrome from *Ldlr*^−*/*−^ and *Gimap6*^−*/*−^ *Ldlr*^−*/*−^ male mice. Mice were maintained on a SFD until 2 months of age, followed by an HFD for 3 months. Dashed boxes highlight 2.7x enlarged regions shown at right, focusing on foam cells (red arrows) and necrotic cores (black arrows). Scale bar, 500 μm. (E) Representative immunofluorescent images of thoracic (top row) and abdominal (bottom row) aortic regions from *Ldlr*^−*/*−^ and *Gimap6*^−*/*−^ *Ldlr*^−*/*−^ male mice from (D), stained for CD68 (macrophages, green), α-smooth muscle actin (α-SMA; smooth muscle cells, magenta), and DAPI (nuclei, blue). Dashed boxes highlight 2x enlarged regions shown at right. The dashed yellow line shows the boundary of the SMC layer. White arrows indicate the infiltration of macrophages. Scale bar, 500 μm. (F) Masson’s trichrome staining of the aortic valve from *Ldlr*^−*/*−^ and *Gimap6*^−*/*−^ *Ldlr*^−*/*−^ mice, showing collagen in blue. Mice were fed an SFD until 2 months of age, then switched to an HFD for 1 month prior to tissue collection. Red arrows indicate foam cell aggregates in the intima; black arrows indicate necrotic cores. Scale bar, 200 µm. (G) Quantification of plaque area and necrotic core area in the aortic valve from (F) using ImageJ. Each dot represents one mouse, bars indicate mean ± SD. Sample sizes: *Ldlr*^−*/*−^ (males, n = 3; females, n = 3); *Gimap6*^−*/*−^ *Ldlr*^−*/*−^ (males, n = 3; females, n = 4). (H) Histological analysis of heart sections from male *Ldlr*^−*/*−^ (top row) and *Gimap6*^−*/*−^ *Ldlr*^−*/*−^ mice (middle and bottom outlined by the dotted box) fed an SFD until 2 months of age, followed by an HFD for 2 months, showing cross-sections of intramyocardial coronary arteries. Columns 1-3: Masson’s trichrome staining shows fibrotic tissue in blue, with asterisks (*) highlighting potential infarcts. Blue arrows indicate normal coronary arteries. Atherosclerotic coronary arteries are illustrated by foam cells (yellow arrows) and necrotic cores (green arrows). Column 4: ORO staining shows lipid-laden foam cells (red) within atherosclerotic coronary arteries (yellow arrows) and their absence in normal arteries (blue arrows). Column 5: immunofluorescence staining for CD68 (macrophages, green), α-smooth muscle actin (α-SMA; smooth muscle cells, magenta), and DAPI (nuclei, blue). For all columns, the bottom row shows 3x magnified views of dashed box regions from the *Gimap6*^−*/*−^ *Ldlr*^−*/*−^ sections depicted in the middle row; scale bars, 200□µm. *P* values were calculated by *t*-test in (G), one-way ANOVA in (C) or log-rank (Mantel-Cox) test in (A). Statistical significance is denoted as not significant (ns), \**P* < 0.05, \*\**P* < 0.01, \*\*\**P* < 0.001, \*\*\*\**P* < 0.0001.

To determine the direct cause of death, we performed a detailed histopathological analysis of the coronary arteries. After only four weeks on an HFD, *Gimap6*^−*/*−^*Ldlr*^−*/*−^ mice developed advanced plaques with foam cells, necrotic cores in both the aortic valve and coronary ostia that were not observed in *Ldlr*^−*/*−^ controls (Figures 2F-G). While the coronary arteries of *Ldlr*^−*/*−^ mice appeared patent, *Gimap6*^−*/*−^*Ldlr*^−*/*−^ mice developed severe and complex atherosclerosis deep within the heart muscle, forming plaques in the smaller intramyocardial branches after two months on an HFD (Figure 2H). Masson’s trichrome staining revealed large, occlusive plaques in the coronary arteries with prominent necrotic cores and perivascular fibrosis extending into the surrounding myocardium, indicative of ischemic damage (Figures 2H1-3). The atherosclerotic lesions were confirmed to be lipid-rich with abundant ORO-positive foam cells (Figure 2H4). Immunofluorescence analysis established that the coronary plaques were heavily infiltrated with CD68^+^ macrophages, which disrupted smooth muscle cell continuity in the tunica media (Figure 2H5).

Taken together, these mouse data show that loss of GIMAP6 is sufficient to cause spontaneous atherosclerosis under standard lipid conditions but that its loss also synergizes with *Ldlr* deficiency-associated hyperlipidemia to promote advanced, occlusive coronary atherosclerosis. Importantly, the extensive immune cell infiltration of the media exceeds the cardiovascular pathology observed in classic murine models, suggesting that *Gimap6* loss results in a lethal inflammatory vasculopathy more complex than hyperlipidemia-driven atherosclerosis.

### *Gimap6* suppresses a lethal macrophage foam cell phenotype

To confirm the cell types in which *Gimap6* is expressed, we performed single-cell transcriptomic analyses of mouse cardiovascular tissues. Indeed, *Gimap6* mRNA was highly enriched in endothelial cells and macrophages of the aorta (Figures 3A-B).^22,23^ We hypothesized that GIMAP6 may plays an essential, cell-autonomous role within macrophages in restricting the initiation of atherosclerosis.

**Figure 3.**
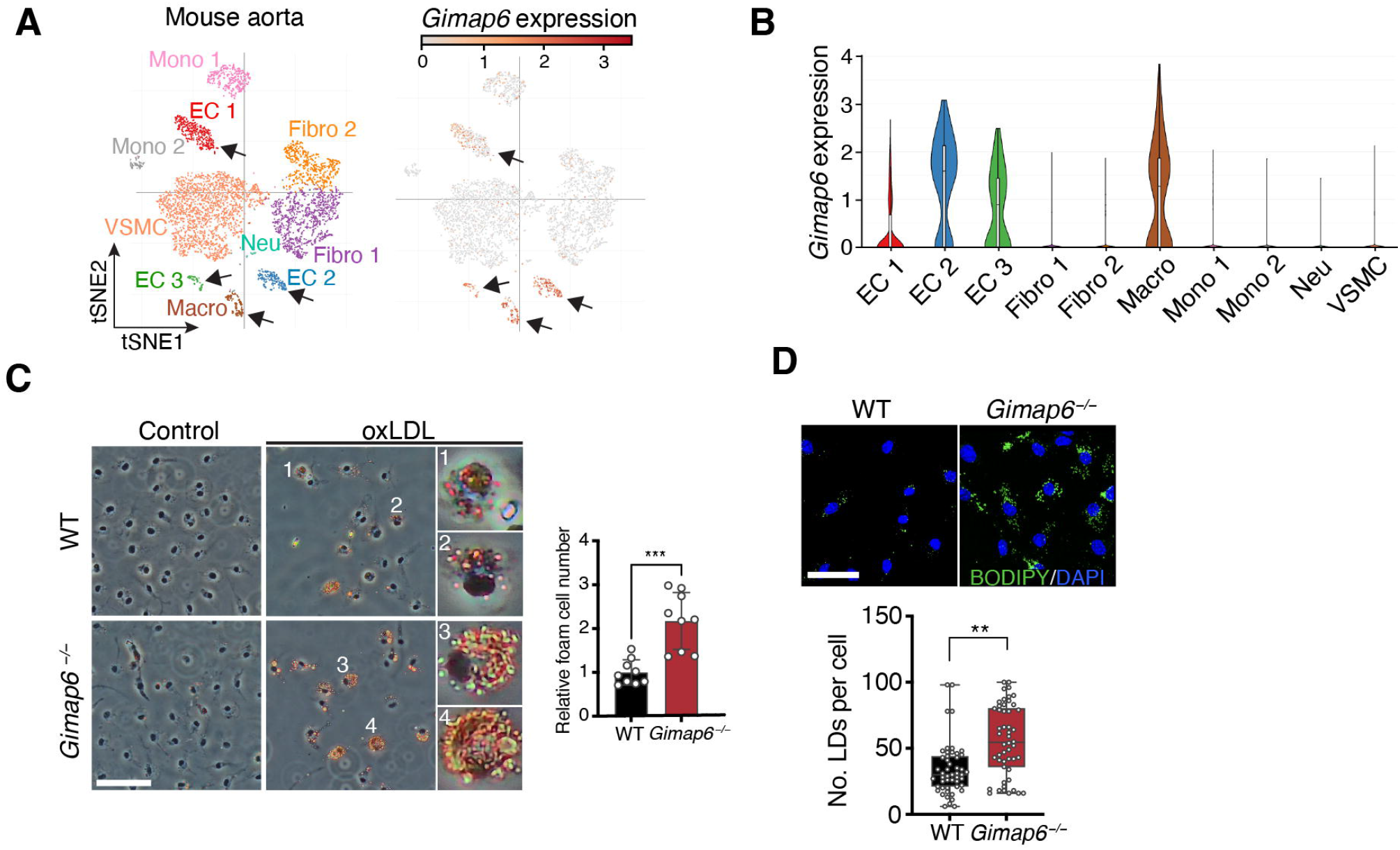
*Gimap6* deficiency promotes foam cell formation. (A) Single-cell transcriptomic analysis of the mouse aorta, obtained from the Single Cell Portal (SCP289)^26^. Left: t-SNE plots display distinct cell clusters, including endothelial cells (EC 1–3), fibroblasts (Fibro 1–2), macrophages (Macro), monocytes (Mono 1–2), neurons (Neu), and vascular smooth muscle cells (VSMC). Right: *Gimap6* expression across these clusters, with color intensity representing normalized expression levels. Black arrows show clusters with high *Gimap6* mRNA expression. (B) Violin plot of *Gimap6* expression across the cell clusters identified in (A). (C) Oil Red Oil staining of bone marrow-derived macrophages (BMDMs) from WT and *Gimap6*^−*/*−^ mice treated with 50 μg/mL oxLDL for 24 hours. Lipid droplets are stained in red and foam cells are indicated by red arrows. Insets (1–4) show magnified views of the indicated foam cells. Scale bar, 50 μm. Right: quantification of relative foam cell numbers per field (n = 9–10 fields per experiment). (D) Top: representative images of unstimulated bone marrow-derived macrophages (BMDMs) from WT and *Gimap6*^−*/*−^ mice stained with BODIPY 493/503 (neutral lipids, green) and DAPI (nuclei, blue). Scale bar, 50 μm. Bottom: quantification of the number (No.) of lipid droplets (LDs) per cell, with 50–100 cells analyzed per group. *P* values were calculated by a two-tailed unpaired *t*-test in (C) and (D). Statistical significance is denoted as not significant (ns), \**P* < 0.05, \*\**P* < 0.01, \*\*\**P* < 0.001.

To test this hypothesis, we isolated BMDMs from WT and *Gimap6*^−*/*−^ mice and used oxidized LDL (oxLDL) exposure to induce lipid uptake and foam cell generation.^24^ ORO staining revealed a significant increase in lipid-laden foam cells in *Gimap6*^−*/*−^ BMDMs compared to WT controls (Figure 3C). BODIPY 493/503 fluorescence staining confirmed amplified lipid accumulation in *Gimap6*^−*/*−^ BMDMs, as indicated by the higher number of lipid droplets per cell in standard culture medium (Figure 3D). Thus, GIMAP6-deficient macrophages are intrinsically re-programmed to over-accumulate lipids. Together, these data establish a cell-intrinsic role for GIMAP6 in lowering lipid uptake and foam cell formation, thereby protecting against ASCVD.

### Pathogenic GIMAP6 variants link protein instability to a human vasculopathy without hyperlipidemia

We previously identified three individuals with rare homozygous stop-gain or missense hypomorphic variants Trp86Ter (W86*) and Gly153Val (G153V).^9,25^ The W86* variant occurred in two children from Gaza who were lost to follow-up. The patient carrying the G153V variant (Pt. 1) suffered from biopsy-confirmed inflammatory vasculopathy and acute focal ischemic lesions in her 20s, and later developed a borderline prolonged QTc interval and mild cardiomegaly (not shown), but has not yet suffered a large ischemic event.^9^ Crucially, a longitudinal analysis of her clinical data revealed a normal plasma lipid profile, including low LDL and cholesterol with mildly elevated triglycerides (not shown), which is consistent with key cardiac and lipid phenotypes observed in *Gimap6*-deficient mice. In searching for additional GIMAP6-deficient patients, we identified from immunodeficiency clinical cohorts two new patients who carried rare, homozygous missense variants in GIMAP6 (Figure 4A). Pt. 2 carrying the Ile44Thr variant (I44T; 7-150325555-A>G; CADD 22.8; AlphaMissense classification likely pathogenic; minor allele frequency = 0.0053 in South Asia) presented with deep venous thrombosis and a strong family history of midlife stroke, myocardial infarction (MI), and CAD (Figure 4A). While the patient has not yet suffered a major ischemic event, a recent echocardiogram at age 38 showed preserved systolic function (ejection fraction 64%) but severe diastolic dysfunction (Mitral E/A ratio of 2.68), and her electrocardiogram displayed nonspecific T-wave abnormalities consistent with myocardial ischemia (Figure 4A). These findings suggest early evidence of GIMAP6-driven vasculopathy. Triglycerides were moderately elevated, and cholesterol and LDL were only slightly above the normal range indicating no severe lipidemia (not shown). Pt. 3, a 10-year-old boy from a consanguineous Persian family with the homozygous Val126Met variant (V126M; 7-150628222-C-T; CADD 13.38; AlphaMissense classification likely benign; minor allele frequency = 0.0013 in non-Finnish Europeans), presented at age 9 with an acute-onset stroke after a childhood history of recurrent infections and immune dysregulation. Brain MRI and angiography revealed multiple acute infarcts and diffuse cerebral vasculitis, with multifocal narrowing of major cerebral arteries. Importantly, this severe vascular event occurred in the context of a normal lipid profile (total cholesterol: 157 mg/dL, LDL: 81 mg/dL) and was accompanied by markedly elevated systemic inflammatory markers, including CRP and TNFα. Despite aggressive immunomodulatory and intensive critical care, the patient rapidly deteriorated and died from massive cerebral edema and herniation (Figure 4A).

**Figure 4.**
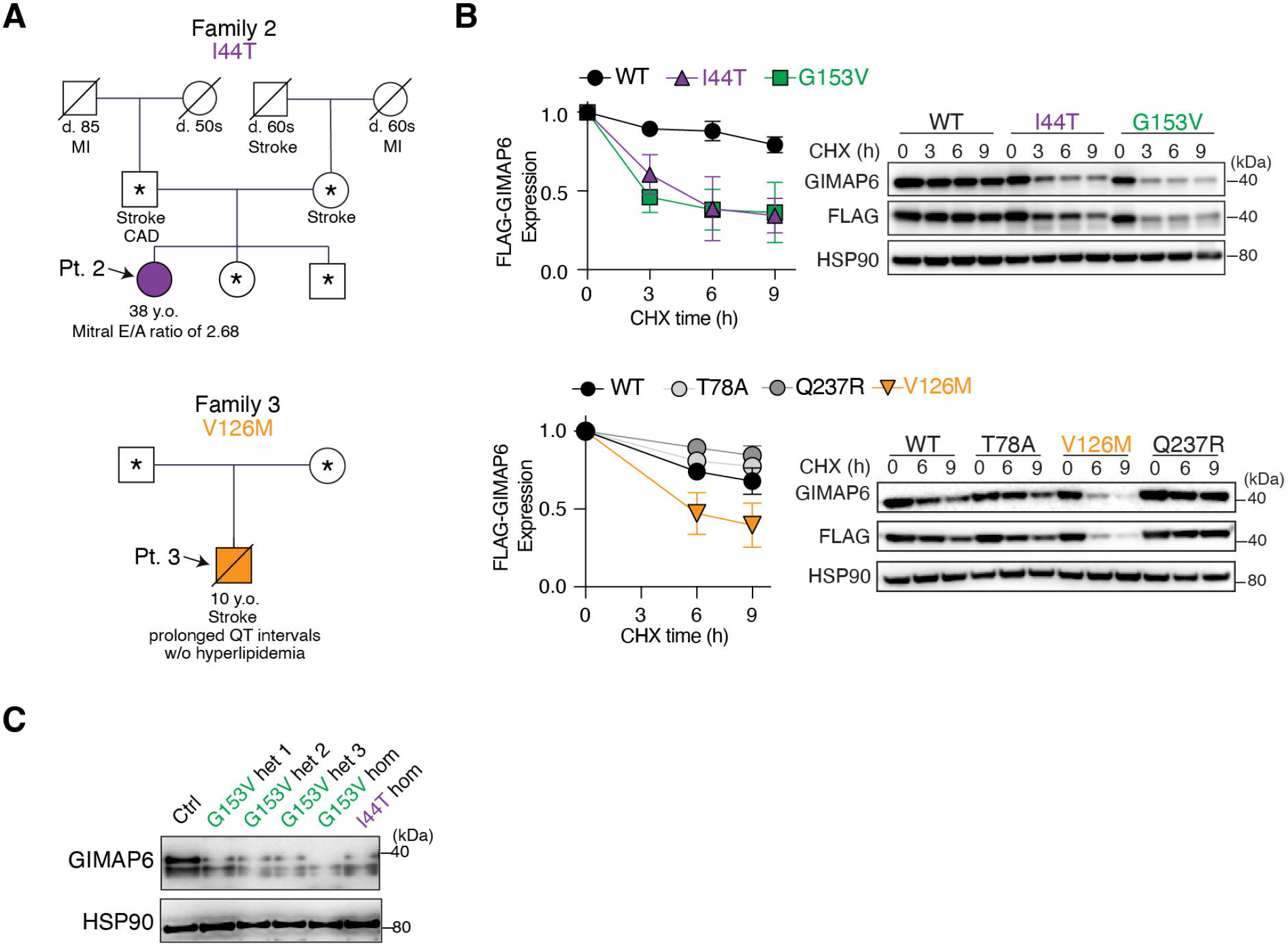
Pathogenic *GIMAP6* variants code unstable proteins results in vasculopathy in human. (A) Pedigrees of two families carrying pathogenic *GIMAP6* variants. Affected individuals are shaded (circles, female; squares, male). The proband in each family is marked by an arrow. Asterisk (*) indicates individuals without available sequencing data. Myocardial infarction, MI; years old, y.o. (B) Left: Densitometric quantification plots of GIMAP6 expression normalized to the 0-hour time point and HSP90 to quantify the protein degradation ratio. Right: representative immunoblots showing protein degradation of FLAG-tagged GIMAP6 over time in a cycloheximide (CHX) chase assay performed with HEK293T cells transiently transfected with matched vectors encoding WT, I44T, G153V, T78A, V126M, or Q237R GIMAP6. (C) Representative immunoblots of GIMAP6 and HSP90 using peripheral blood mononuclear cell lysates from I44T and G153V homozygotes (hom), G153V heterozygous (het 1–3) family members, and healthy donors (ctrl). *P* values were calculated by two-way ANOVA for (B). Statistical significance is indicated as not significant (ns), \*\**P* < 0.01.

We suspected that the I44T and V126M mutations in Pts. 2 and 3, respectively, decreased protein stability, similar to what we found for the G153V variant protein in Pt. 1 in our previous study.^9^ Therefore, we performed cycloheximide (CHX) chase experiments, which revealed that all three variants showed significantly faster degradation of the GIMAP6 protein compared to wild-type (WT) or common variants Gln237Arg (Q237R) and Thr78Ala (T78A) (Figure 4B). Immunoblotting revealed that GIMAP6 protein expression was also markedly decreased in lysates from peripheral blood mononuclear cells (PBMC) from Pts. 1 and 2, compared to a healthy normal donor (Figure 4C). These results establish that these patients with severe vascular phenotypes carry hypomorphic GIMAP6 variants.

Collectively, our data from three unrelated families support the concept that rare hypomorphic GIMAP6 variants cause protein instability, which drives severe, normolipidemic vasculopathy in humans.

## DISCUSSION

The foundation of our understanding of ASCVD is that elevated plasma cholesterol and lipoproteins drive cholesterol into macrophages that become pathogenic inflammatory mediators aggregated in plaques in the walls of critical arteries. The underlying pathogenetic mechanism that creates the mass-action drive for cellular cholesterol accumulation is the failure of LDL uptake and metabolism due to reduced LDLR expression in the liver either due to genetic deficiencies or natural aging-related down-regulation. Here we examine the situation in which circulating cholesterol and lipoproteins are normal in the absence or diminished expression of GIMAP6, but lipid uptake increases in macrophages. This reveals an independent mechanism that causes fatal ASCVD on a standard fat diet and synergizes to exacerbate disease caused by *Ldlr* deficiency in mice. Hence, GIMAP6 is a key regulator of cardiovascular homeostasis, a function that extends beyond the canonical immunological roles of the GIMAP family. We observe that GIMAP6 deficiency leads to ASCVD in normolipidemia by changing the lipid uptake in macrophages.

Our new understanding of the GIMAP6 as a fundamental driver of ASCVD suggests new approaches for therapeutic and ASCVD risk evaluation. Our data suggest that GIMAP6 could be a potential therapeutic target. By targeting GIMAP6 itself would carry a lower risk of systemic off-target effects, as its tissue expression is restricted primarily to hematopoietic and endothelial cells and absent from parenchymal cells in the heart and other vital organs. The development of small molecule chaperones or genetic strategies to augment GIMAP6 activity directly or its downstream pathway represents a novel therapeutic paradigm for this disease. Targeting this pathway in macrophages could offer specific treatment for a previously unrecognized high-risk population.

## RESOURCE AVAILABILITY

### Lead contact

Requests for further information and resources should be directed to and will be fulfilled by the lead contact, Michael J. Lenardo.

### Materials availability

Materials generated in this study will be made available upon reasonable request from the lead contact.

### Data and code availability

- The human exome and genome sequencing data reported in this paper have been deposited at dbGaP, which is a restricted access database for qualified investigators.
- Any additional information required to reanalyze the data reported in this paper is available from the lead contact upon request.
- Data are publicly available as of the date of publication. These accession numbers for the datasets are listed in the key resources table.

## ACKNOWLEDGMENTS

We also thank Madeleine Cule, Divyanshi Srivastava, Joanne Berghout and Bioinformatics and Computational Biosciences Branch (BCBB) for their bioinformatics expertise and technical support. Margaret Abaandou for regulatory support, Wesley Tung for assistance with plasmid construction, and Drs. Alessandra Brofferio and Ahmet Ozen for clinical assistance. We thank Dr. Peter Grayson, Ronald N. Germain, and Pamela L. Schwartzberg for critically reading the manuscript. Some figures were created using BioRender.com. This paper is dedicated to the memory of Guido D. Lenardo, M.D.

This research was supported [in part] by the Intramural Research Program of the National Institutes of Health (NIH). The contributions of the NIH author(s) are considered Works of the United States Government. The findings and conclusions presented in this paper are those of the author(s) and do not necessarily reflect the views of the NIH or the U.S. Department of Health and Human Services.

## AUTHOR CONTRIBUTIONS

Conceptualization, C.X., L.Z., M.J.L.; Methodology, C.X., D.S., Z.H.Y., E.E., Y.Z., S.W., R.V., D.C., S.M.H., C.S.Z., A.T.R., H.C.S., L.Z., M.J.L.; Investigation, C.X., K.A., D.S., S.N., P.D.J., J.C., M.H.Z., A.Y.P., J.T., M.S., H.J., Q.L., E.E., J.K., R.V., L.Z.; Visualization, C.X., K.A., D.S., S.N., M.S., Q.L., R.V.; Funding Acquisition, H.C.S., M.J.L.; Project Administration, C.X., L.Z., M.J.L.; Supervision, H.C.S., L.Z., M.J.L.; Writing – Original Draft, C.X., K.A., M.J.L.; Writing – Review & Editing, C.X., K.A., H.C.S., L.Z., and M.J.L.

## DECLARATION OF GENERATIVE AI AND AI-ASSISTED TECHNOLOGIES

During the preparation of this work, the author(s) utilized artificial intelligence-based language models (ChatGPT 4o and Gemini Pro 2.5) to assist with language editing and coding. The purpose was to identify and correct grammatical errors, spelling, and punctuation, and to improve the overall readability of the text. After using these tools, the author(s) reviewed and edited the content as needed and take full responsibility for the content of the publication.

## EXPERIMENTAL MODEL AND STUDY PARTICIPANT DETAILS

### Human participants

Written informed consent was obtained from all participants in accordance with local ethics and Institutional Review Board (IRB) guidelines. Patient 1 was enrolled in NIH protocol NCT00246857, and genetic sequencing was performed as previously reported. Patient 2 was enrolled in NIH protocol NCT00018044, with clinical and laboratory samples collected during inpatient visits. Patient 3 was enrolled under protocol IR.TUMS.CHMC.REC.1398.030, approved by the Tehran University of Medical Sciences Ethics Committee. Control samples and healthy donor blood were obtained through separate IRB-approved protocols (NCT00128973) at the NIH Clinical Center. Detailed clinical summaries, laboratory findings, and family histories for all subjects are provided in the Supplementary Information.

### Mouse strains, housing, and diet

All animal procedures were approved by the NIAID Animal Care and Use Committee (protocol LISB-7E) and followed NIH and Department of Health and Human Services guidelines. B6N(Cg)-*Gimap6*<tm1b(KOMP)Wtsi>/J mice were generated by the Knockout Mouse Project (KOMP) and maintained as previously described.^9^ B6.129S7-*Ldlr*<tm1Her>/J mice (JAX Stock No. 002207, obtained from Taconic Biosciences) were crossed with *Gimap6*-deficient mice to generate *Gimap6*^−/−^ *Ldlr*^−/−^ double knockout animals. Experimental cohorts were derived from intercrosses of *Gimap6*^+/−^, *Gimap6*^+/−^ *Cd36*^−/−^ or *Gimap6*^+/−^ *Ldlr*^−/−^ heterozygous breeders. Littermate knockouts and controls were co-housed to minimize cage and microbiota effects. Mice were housed under specific pathogen-free (SPF) conditions (22□±□2°C, 12□h light/dark cycle, 40–60% humidity) with up to five per individually ventilated cage, provided autoclaved chow and water ad libitum, and environmental enrichment. For survival studies, mice were maintained on standard chow. For atherosclerosis experiments, 8-week-old mice were placed on a high-fat diet (40% kcal fat, 1.25% cholesterol; Research Diets, #D12108C) for the indicated durations. Animals were allowed to age naturally unless removed by veterinary staff due to non-age-related health concerns, in which case they were euthanized by CO_2_ inhalation followed by cervical dislocation.

### Alternative PCR genotyping

B6N(Cg)-*Gimap6*<tm1b(KOMP)Wtsi>/J mice were genotyped using allele-specific PCR to distinguish wild-type (WT) and knockout (KO) alleles. The KO allele (617□bp) was amplified using primers: Forward 5′-GCT ACC ATT ACC AGT TGG TCT GGT GTC-3′ and Reverse 5′-ATA TAA AGA CCC TCC CTT CAC CGC C-3′. The WT allele (415□bp) was amplified using primers: Forward 5′-TTG CTG CCA TCT TCC TTA CAA GTG C-3′ and Reverse 5′-AAG TTA TTC CAC ACG TTT GGG GG-3′. PCR was performed in 50□µl reactions containing 25□µl 2× MyTaq™ Red Mix (Bioline), 0.1□µM of each primer, 1□µl genomic DNA, and nuclease-free water. Thermocycling conditions were as follows: 94°C for 5□min; 10 touchdown cycles of 94°C for 15□s, 65→55°C for 30□s (–1°C/cycle), and 72°C for 40□s; then 25 cycles of 94°C for 15□s, 55°C for 30□s, and 72°C for 40□s; with a final extension at 72°C for 10□min. PCR products were separated by agarose gel electrophoresis (KO = 617□bp; WT = 415□bp). B6.129S7-*Ldlr*<tm1Her>/J mice (JAX Stock No. 002207) were genotyped according to the Jackson Laboratory’s standard protocol (Protocol 27075, version 1.2).

### Cell culture

THP-1 cells (ATCC, TIB-202), a human monocytic leukemia line, were cultured in RPMI-1640 medium (Gibco, Thermo Fisher Scientific) supplemented with 10% heat-inactivated fetal bovine serum (FBS; Gibco), 2□mM L-glutamine, 100□IU/mL penicillin, and 100□μg/mL streptomycin. Cells were maintained at 37°C in a humidified 5% CO_2_ incubator at 2×10□ to 6×10□ cells/mL and passaged every 2–3 days to ensure exponential growth and > 90% viability (trypan blue exclusion). HEK293T cells (ATCC, #CRL-3216) were cultured in Dulbecco’s Modified Eagle Medium (DMEM; Gibco, Thermo Fisher Scientific) supplemented with 10% fetal bovine serum and 1% penicillin-streptomycin and maintained in 100□mm tissue culture plates at 70–90% confluency. Primary human CD14^+^ monocytes were isolated from fresh peripheral blood mononuclear cells (PBMCs) collected from healthy adult donors. PBMCs were prepared using Ficoll-Paque™ Plus (Cytiva, #17144002) density gradient centrifugation, followed by positive selection with CD14 MicroBeads (Miltenyi Biotec, #130-050-201) and magnetic-activated cell sorting (MACS) per manufacturer’s instructions. The resulting monocyte population was ≥95% pure, as confirmed by flow cytometry. Purified CD14^+^ monocytes were cultured in RPMI-1640 medium supplemented as described above, seeded at 1×10□ cells/mL, and used within 48□h to preserve monocyte identity and function.

### Plasmid mutagenesis

mCherry-T2A-FLAG-GIMAP6 plasmids (wild-type and mutant variants) were synthesized using the Gene Synthesis service from GenScript. Plasmid maps for all constructs are available upon request.

### Survival analysis

To assess the impact of *Gimap6* deficiency on survival, Kaplan-Meier analysis was performed in mice maintained on standard chow. *Gimap6*^−/−^, *Gimap6*^+/−^, and *Gimap6*^+/+^ mice were generated through intercrosses of *Gimap6*^+/−^ heterozygotes. Similarly, *Gimap6*^−/−^ *Ldlr*^−/−^, *Gimap6*^+/−^ *Ldlr*^−/−^, and *Ldlr*^−/−^ mice were produced from controlled heterozygous matings. Breeding was continuous, and mice were enrolled in survival cohorts at weaning (postnatal day 21), with heterozygous female *Gimap6*^+/−^ breeder mice included in the analyses. Age- and sex-matched littermates from different litters were co-housed (up to five per cage) under identical SPF conditions to minimize environmental and microbiota variability. Mice were monitored weekly and euthanized at predefined humane endpoints per institutional guidelines. Survival data were pooled across multiple litters and generations, and Kaplan-Meier curves were generated and analyzed using GraphPad Prism v10.

### Body weight measurements

Body weight was recorded weekly for all mice enrolled in the survival cohorts at the indicated time points using a calibrated digital scale. *Gimap6*^−/−^, *Gimap6*^+/−^, and *Gimap6*^+/+^ mice were age- and sex-matched, housed under SPF conditions with ad libitum autoclaved standard chow and water (15 per genotype and sex; 3 to 5 per group, pooled from three independent experiments). Data are presented as mean ± standard deviation (SD).

### Heart weight measurements

Gross heart morphology and heart weight were assessed in *Gimap6*^−/−^, *Gimap6*^+/−^, and *Gimap6*^+/+^ mice maintained on standard chow at the indicated time points. After euthanasia, hearts were excised, rinsed in cold PBS, blotted dry, and immediately photographed under standardized lighting and magnification to document gross anatomy. Hearts were weighed using a calibrated analytical balance. To minimize animal use, age- and sex-matched controls were selected to match shorter-lived knockout mice, where applicable. For each genotype, sex, and age group, 12 mice were analyzed (3 to 5 per group, pooled from three independent experiments). Results are presented as mean ± SD.

### Cardiac troponin quantification

Plasma cardiac troponin I (cTnI) was measured using a mouse-specific ELISA kit (MyBioSource, #MBS766175; sensitivity 4.7□pg/mL; dynamic range 7.8–500□pg/mL) following the manufacturer’s instructions. Plasma (50□µL per well) was read at 450□nm on a VersaMax Microplate Reader (Molecular Devices, #VERSAMAX), and cTnI concentrations were calculated from a recombinant protein standard curve. For time-course analysis, ∼150□µL whole blood was collected every 4 weeks (8 to 24 weeks of age; n = 3 to 7 mice per time point, pooled from three independent experiments) via tail vein into lithium heparin tubes (SARSTEDT, #41.1393.105). Blood from moribund mice (per veterinary-defined humane endpoints) was collected by cardiac puncture under CO_2_ anesthesia; age- and sex-matched controls were euthanized in parallel. Plasma was isolated by centrifugation (3,000 × g, 10□min, 4°C), aliquoted, and stored at −80°C until analysis.

### Mouse histopathology

For paraffin sections, tissues were fixed overnight at 4°C in 10% neutral buffered formalin (Sigma-Aldrich, #HT5012), transferred to 80% ethanol, processed on a Tissue-Tek VIP system (Sakura, GMI Inc.), embedded in paraffin, sectioned at 5□μm (rotary microtome), and mounted on Superfrost Plus slides (Thermo Fisher, #12-550-15). Slides were dried overnight at 37°C before staining. For frozen sections, tissues were embedded in Tissue-Tek® O.C.T. (Sakura, #4583), snap-frozen in isopentane over liquid nitrogen, stored at −80°C, sectioned at 5□μm (cryostat), mounted, air-dried at room temperature, and either stained immediately or stored at −80°C.

### 2,3,5-triphenyltetrazolium chloride staining

Myocardial viability was assessed by 2,3,5-triphenyltetrazolium chloride (TTC) staining of freshly excised mouse hearts. *Gimap6*^−*/*−^ mice, which typically appear acutely ill and die within ∼24□h after symptom onset, were euthanized at moribund stages alongside age- and sex-matched controls. Immediately post-euthanasia, hearts were rinsed in cold PBS (pH 7.4; Gibco, #10010023), transversely sliced into 4 to 5 sections, and incubated in 1% TTC solution (Sigma-Aldrich, #T8877) freshly dissolved in PBS at 37°C for 50□min in the dark (CO_2_ incubator; Thermo Fisher, Heracell™ 150i/240i). After staining, slices were fixed in 10% neutral buffered formalin (Sigma-Aldrich, #HT5012) at room temperature for at least 24□h. Images were acquired using a digital camara in ambient lighting. Viable myocardium stained red (formazan), while infarcted regions remained pale.

### Masson’s trichrome staining

Cardiac fibrosis and extracellular matrix were assessed on formalin-fixed, paraffin-embedded (FFPE) heart sections (5□μm) using the Masson’s Trichrome Stain Kit (VitroView, #VB-3016). Sections were deparaffinized in xylene (2 × 6□min), rehydrated through graded ethanol (100%, 95%, 70%), and rinsed in distilled water. Slides were mordanted in preheated Bouin’s solution (60°C, 1□h), cooled, and rinsed in tap and distilled water. Nuclear staining was performed with freshly prepared Weigert’s working hematoxylin (10□min), followed by Biebrich Scarlet–Acid Fuchsin cytoplasmic staining (5□min). Sections were treated with phosphomolybdic–phosphotungstic acid (10□min), stained with Aniline Blue (10□min), dipped briefly in 1% acetic acid, and rinsed. Slides were dehydrated through 95% and 100% ethanol, cleared in xylene (3×5□min), and mounted with Permount (Fisher Scientific, #SP15-100). Collagen stained blue, muscle fibers red, and nuclei black.

### Echocardiography (M-mode and Doppler)

Cardiac structure and function were assessed by high-resolution transthoracic echocardiography. Mice were lightly anesthetized with 1 to 1.5% isoflurane via nose cone and placed supine on a heated rail-mounted platform with ECG leads and rectal temperature probe (VisualSonics, Toronto, Canada). Heart rate, respiratory rate, and core temperature were continuously monitored and maintained within physiological limits. Chest fur was removed by shaving and depilatory cream; ultrasound gel was applied for acoustic coupling. Imaging was performed using a VevoF2 imaging system (FUJIFILM VisualSonics) with a UHF46X linear transducer with a bandwidth of 46-20 mHz (FUJIFILM VisualSonics). Two-dimensional (2D) and M-mode images were acquired from parasternal long-axis and mid-papillary short-axis views. M-mode tracings measured LV internal diameter at end-diastole (LVIDd) and end-systole (LVIDs), with ejection fraction (EF) calculated as EF (%) = [(LVIDd − LVIDs) / LVIDd] × 100. Pulsed-wave Doppler imaging assessed ascending aortic outflow and valve function; additional parameters (e.g., vessel diameter, pulse wave velocity) were acquired as needed. Each session lasted ∼15□min per animal. Mice recovered on a warming pad or were euthanized per protocol. Quantitative measurements are provided in Tables S1–2.

### ECG transmitter implantation and analysis

Long-term cardiac monitoring was performed using surgically implanted ECG telemetry transmitters (Data Sciences International, ETA-F10) under sterile conditions. Mice were anesthetized with 2% isoflurane and positioned in dorsal recumbency. The body temperature was maintained at 37□°C using a water blanket and monitored with rectal temperature probe. Preoperative analgesia (Buprenorphine ER®, 1□mg/kg subcutaneously) was administered. The chest and abdomen were shaved and disinfected with alternating chlorhexidine and 70% ethanol scrubs. A midline abdominal incision was made caudal to the xiphoid process, and the transmitter body was implanted in the lower abdominal cavity and secured with non-absorbable 6-0. The ECG leads were passed from the abdominal cavity to the exterior and tunneled subcutaneously: the positive lead to the lower chest pectoral muscles, the negative lead to the upper right pectoral muscles. Both leads and the abdomen were anchored with non-absorbable 6-0 sutures. The skin incisions were closed with wound clips; Bupivacaine was applied topically. One milliliter of saline was given by subcutaneously on back. Postoperative care included oxygenated warmed recovery, housing in oxygen-enriched ICU cages on heated water blankets for 3 days, and twice-daily monitoring (Murine Phenotyping Core or surgical core staff) for pain, dehydration, wound healing, and food intake. Analgesia (Buprenorphine ER®, 1□mg/kg every 2 to 3 days; Meloxicam, 5□mg/kg daily) and subcutaneous fluids (1 to 2□mL as needed) were provided per NHLBI protocols. Staples or wound clips were removed 10 to 14 days post-surgery if incisions were fully healed; mice were euthanized if distress or complications were observed. The ECG data were recorded and analyzed using Ponemah software (Data Sciences International).

### PowerLab ECG analysis

ECG signals were acquired using PowerLab ECG equipment (ADInstruments, Octal Bio Amp). Mice were anesthetized with 2% and positioned in dorsal recumbency. The body temperature was maintained at 37□°C using a heat lamp and monitored with rectal temperature probe. For surface ECG, 29-gauge needle electrodes were inserted subcutaneously into limbs for 4-lead configurations (Lead I and II). Signals were recorded for 2 min per animal using LabChart 8 software (ADInstruments), and Lead I was used for the analysis.

### Oil Red O staining of en face aortae

Mice were euthanized per institutional guidelines, and the full aorta (aortic root to iliac bifurcation) was dissected under a stereomicroscope, rinsed in cold PBS (pH 7.4), and fixed in 10% neutral buffered formalin (Sigma-Aldrich, #HT5012) for 5□min at room temperature. After rinsing in PBS, aortae were stored in sterile saline or distilled water in 6-well plates to prevent dehydration. Oil Red O working solution was freshly prepared by diluting 0.5% (w/v) Oil Red O stock in isopropanol (Abcam, #ab146295) at 3:2 with distilled water, then filtered (Steriflip®, Millipore, #SE1M003M00). Aortae were stained for 30□min at room temperature with gentle agitation, washed in 60% isopropanol (30□min), rinsed in distilled water, and stored in PBS at 4°C. Adventitial fat was removed using micro-dissection scissors and forceps, and aortae were opened longitudinally along the ventral side. For en face imaging, stained aortae were mounted between two glass slides: a drop of distilled water was placed on a Superfrost Plus slide (Thermo Fisher, #12-550-15), the aorta (adventitia side down) was flattened with Extra Fine Graefe Forceps (Fine Science Tools, #11150-10), and a second slide was placed on top. Corners were sealed with Loctite Waterproof Sealant (#908570); after drying (∼30 to 60□min), the sandwich was placed vertically in a water-filled dish to promote capillary action. Air bubbles were removed with a 31G insulin needle, and remaining edges were sealed. Slides were air-dried (1□h), stored at 4°C in light-protected boxes, and imaged within one week. Digital microscope images were captured at matched magnification and exposure settings for the aortic arch and thoracoabdominal regions.

Lipid deposition was quantified using ImageJ software (NIH, ImageJ bundled with Java 8) as previously described.^27^ Images were converted to 8-bit grayscale, thresholded to isolate Oil Red O–positive areas and analyzed using the polygon tool to calculate: Percent lesion area = (Oil Red O–positive area / total aortic area) × 100. All images were analyzed independently under identical thresholding parameters, blinded, by at least two investigators.

### Histology and immunofluorescence

FFPE heart and aorta sections were mounted on Superfrost Plus slides (Thermo Fisher, #12-550-15), deparaffinized in xylene (2 × 6□min), rehydrated through graded ethanol (100%, 95%, 70%), and rinsed in distilled water. Antigen retrieval was performed in citrate buffer (10□mM sodium citrate, pH 6.0; BioLegend, #828502) at 95 to 100°C for 15 to 20□min, then cooled at room temperature for 30□min. After PBS washes, sections were blocked for 1□h at room temperature in 5% normal goat serum, 0.3% Triton X-100 in PBS. Primary antibodies (anti-CD68, E3O7V, Cell Signaling, #97778, 1:200; anti-α-smooth muscle actin, R&D Systems, #MAB1420, 10□µg/mL) were diluted in 1% BSA, 0.3% Triton X-100 in PBS and incubated overnight at 4°C. The next day, sections were washed (PBS + 0.3% Triton X-100) and incubated for 1□h at room temperature in the dark with secondary antibodies: goat anti-mouse IgG (H+L) Alexa Fluor Plus 488 (Thermo Fisher, #A32723; RRID: AB_2633275) or goat anti-rabbit IgG (H+L) Alexa Fluor Plus 647 (Thermo Fisher, #A32733; RRID: AB_2633282). Nuclei were counterstained with DAPI (GLPBIO, #GC43378), and slides were mounted with Fluoromount-G (SouthernBiotech, #0100-01) and stored at 4°C protected from light. Fluorescence imaging was performed on a Leica SP8 confocal microscope under identical exposure settings, and image analysis was conducted using Imaris software (v10.1.1; Oxford Instruments).

### Immunoblot

Mouse spleens, bone marrow–derived macrophages (BMDMs), THP-1 cells, and human peripheral blood mononuclear cells (PBMCs) were lysed in RIPA Lysis and Extraction Buffer (Thermo Fisher Scientific, #89901) supplemented with Halt Protease and Phosphatase Inhibitor Cocktails (Thermo Fisher Scientific, #78439 and #78427). Lysates were incubated on ice for 30 minutes and centrifuged at 14,000 rpm for 10 min at 4□°C. Protein concentrations were determined using the BCA Protein Assay Kit (Thermo Fisher Scientific, #23225). Equal amounts of total protein (25□μg per lane) were mixed with NuPAGE LDS Sample Buffer (4×) (Thermo Fisher Scientific, #NP0008) and Reducing Agent (10×) (Thermo Fisher Scientific, #NP0004), heated at 70□°C for 10 min, and resolved by electrophoresis on NuPAGE 4–12% Bis-Tris Protein Gels (Thermo Fisher Scientific, #NP0336BOX). Proteins were transferred onto nitrocellulose membranes (Bio-Rad, #1620112) using standard wet transfer methods. Membranes were blocked with EveryBlot Blocking Buffer (Bio-Rad, #12010020) for 1 h at room temperature, then incubated overnight at 4□°C with primary antibodies diluted in blocking buffer. Primary antibodies included anti-human GIMAP6 (MAC445) ^28^, anti-mouse GIMAP6 (MAC431, MAC432, MAC434, or a mixture of all three) ^28^, anti-CD68 (Cell Signaling Technology, #97778S), anti-β-actin (BioLegend, #664802), and anti-Hsp90 (BioLegend, #675402). After washing in TBST (0.05% Tween-20), membranes were incubated for 1 h at room temperature with HRP-conjugated secondary antibodies (SouthernBiotech, 1:2,000 dilution). Protein bands were visualized using chemiluminescent substrates, including Immobilon Classico or Forte Western HRP Substrate (Millipore, #WBLUC0500 and #WBLUF0500), SuperSignal West Femto Maximum Sensitivity Substrate (Thermo Fisher Scientific, #34096), or SuperSignal West Dura Extended Duration Substrate (Thermo Fisher Scientific, #34076). For membrane stripping and reprobing, membranes were treated with Restore Western Blot Stripping Buffer (Thermo Fisher Scientific, #21063) for 1 h at room temperature, followed by three 5-min washes in TBST. Stripping efficiency was verified by incubating membranes with secondary antibody alone and imaging on an Azure 500 system (Azure Biosystems).

### Transfection and cycloheximide chase assay

HEK293T cells were seeded in DMEM with 10% heat-inactivated FBS and 1% penicillin-streptomycin in 12-well plates and grown to ∼70% confluency. Plasmids were diluted in Opti-MEM™ Reduced Serum Medium (Thermo Fisher) and transfected using Lipofectamine™ 3000 (Thermo Fisher, #L3000075) per manufacturer’s instructions. Cells were incubated at 37°C in a humidified CO_2_ incubator. After 24□h, transfected cells were treated with cycloheximide (50□μg/mL; GlpBio, #GC17198) and lysed at 0, 3, 6, or 9□h for Western blot analysis. FLAG-GIMAP6 expression was quantified relative to time 0 using ImageJ and normalized to HSP90.

### Bone marrow–derived macrophage isolation and culture

Bone marrow-derived macrophages (BMDMs) were isolated and cultured as previously described ^29^, with minor modifications□. Mice were euthanized, and femurs were harvested. Bone marrow was flushed using a 25G needle attached to a 10□mL syringe filled with cold sterile PBS, collected in 50□mL conical tubes on ice, and centrifuged at 500×g for 10□min at room temperature. Pellets were resuspended in macrophage complete medium (DMEM + 20□ng/mL recombinant murine M-CSF; GenScript, #Z02930), filtered through a 70□µm cell strainer (Corning) to remove debris, and counted by hemocytometer. Cells were seeded at 4×10□ per 10□cm non-tissue culture-treated Petri dish in 10□mL medium and maintained at 37°C, 5% CO_2_. On day 3, 5□mL of fresh medium was added. On day 7, nonadherent cells were removed by PBS wash, and adherent macrophages were detached with 0.25% Trypsin-EDTA + phenol red (1□mL/dish; Gibco, #25200056) for 3□min at room temperature, yielding typically > 90% macrophages based on morphology and marker expression.

### scRNA-seq

Publicly available single-cell RNA sequencing (scRNA-seq) datasets from mouse heart and aorta were obtained from the Broad Institute’s Single Cell Portal ^26^ .Datasets were selected based on tissue relevance and the availability of processed gene expression matrices and cell-type annotations. Normalized gene expression values and corresponding cell-type labels were downloaded directly from the portal. All visualizations, including t-distributed stochastic neighbor embedding (t-SNE) plots and violin plots, were generated using the portal’s built-in analysis tools. Expression of *Gimap6* (mouse) or *GIMAP6* (human) was queried across annotated cell populations. No additional filtering, normalization, or reclustering was performed; all analyses reflect the original processing pipelines provided with each dataset. The datasets analyzed included human thoracic aorta (SCP1265)^30^, mouse aorta (SCP289)^23^, and mouse heart (SCP283)^22^.

## REFERENCES

1. Roth, G.A., Mensah, G.A., Johnson, C.O., Addolorato, G., Ammirati, E., Baddour, L.M., Barengo, N.C., Beaton, A.Z., Benjamin, E.J., and Benziger, C.P. (2020). Global burden of cardiovascular diseases and risk factors, 1990–2019: update from the GBD 2019 study. Journal of the American college of cardiology 76, 2982–3021.

2. Ross, R. (1999). Atherosclerosis—an inflammatory disease. New England journal of medicine 340, 115–126.

3. Libby, P. (2021). The changing landscape of atherosclerosis. Nature 592, 524–533.

4. Ference, B.A., Ginsberg, H.N., Graham, I., Ray, K.K., Packard, C.J., Bruckert, E., Hegele, R.A., Krauss, R.M., Raal, F.J., Schunkert, H., et al. (2017). Low-density lipoproteins cause atherosclerotic cardiovascular disease. 1. Evidence from genetic, epidemiologic, and clinical studies. A consensus statement from the European Atherosclerosis Society Consensus Panel. Eur Heart J 38, 2459–2472. 10.1093/eurheartj/ehx144.

5. Musunuru, K., and Kathiresan, S. (2019). Genetics of common, complex coronary artery disease. Cell 177, 132–145.

6. Libby, P., Buring, J.E., Badimon, L., Hansson, G.K., Deanfield, J., Bittencourt, M.S., Tokgözoğlu, L., and Lewis, E.F. (2019). Atherosclerosis. Nature Reviews Disease Primers 5, 56. 10.1038/s41572-019-0106-z.

7. Ridker, P.M., Everett, B.M., Thuren, T., MacFadyen, J.G., Chang, W.H., Ballantyne, C., Fonseca, F., Nicolau, J., Koenig, W., and Anker, S.D. (2017). Antiinflammatory therapy with canakinumab for atherosclerotic disease. New England journal of medicine 377, 1119–1131.

8. Björkegren, J.L.M., and Lusis, A.J. (2022). Atherosclerosis: Recent developments. Cell 185, 1630–1645. 10.1016/j.cell.2022.04.004.

9. Gimbrone, M.A., and García-Cardeña, G. (2016). Endothelial Cell Dysfunction and the Pathobiology of Atherosclerosis. Circulation Research 118, 620–636. 10.1161/CIRCRESAHA.115.306301.

10. Moore, Kathryn J., and Tabas, I. (2011). Macrophages in the Pathogenesis of Atherosclerosis. Cell 145, 341–355. 10.1016/j.cell.2011.04.005.

11. Tabas, I., and Bornfeldt, K.E. (2020). Intracellular and Intercellular Aspects of Macrophage Immunometabolism in Atherosclerosis. Circulation Research 126, 1209–1227. 10.1161/CIRCRESAHA.119.315939.

12. Poli, M.C., Aksentijevich, I., Bousfiha, A.A., Cunningham-Rundles, C., Hambleton, S., Klein, C., Morio, T., Picard, C., Puel, A., Rezaei, N., et al. (2025). Human inborn errors of immunity: 2024 update on the classification from the International Union of Immunological Societies Expert Committee. Journal of Human Immunity 1. 10.70962/jhi.20250003.

13. Xiang, C., Park, A.Y., Weber, S.E., Lenardo, M.J., Ozen, A., and Cui, J. (2025). The impact of genetic immune disorders on infections including COVID-19, inflammatory bowel disease and cancer. Nature Immunology. 10.1038/s41590-025-02225-4.

14. Aragam, K.G., Jiang, T., Goel, A., Kanoni, S., Wolford, B.N., Atri, D.S., Weeks, E.M., Wang, M., Hindy, G., Zhou, W., et al. (2022). Discovery and systematic characterization of risk variants and genes for coronary artery disease in over a million participants. Nature Genetics 54, 1803–1815. 10.1038/s41588-022-01233-6.

15. Maffia, P., Mauro, C., Case, A., and Kemper, C. (2024). Canonical and non-canonical roles of complement in atherosclerosis. Nature Reviews Cardiology 21, 743–761.

16. Zhou, Q., Yang, D., Ombrello, A.K., Zavialov, A.V., Toro, C., Zavialov, A.V., Stone, D.L., Chae, J.J., Rosenzweig, S.D., and Bishop, K. (2014). Early-onset stroke and vasculopathy associated with mutations in ADA2. New England Journal of Medicine 370, 911–920.

17. Limoges, M.-A., Cloutier, M., Nandi, M., Ilangumaran, S., and Ramanathan, S. (2021). The GIMAP Family Proteins: An Incomplete Puzzle. Frontiers in Immunology 12. 10.3389/fimmu.2021.679739.

18. Yao, Y., Du Jiang, P., Chao, B.N., Cagdas, D., Kubo, S., Balasubramaniyam, A., Zhang, Y., Shadur, B., NaserEddin, A., and Folio, L.R. (2022). GIMAP6 regulates autophagy, immune competence, and inflammation in mice and humans. Journal of Experimental Medicine 219.

19. Park, A.Y., Leney-Greene, M., Lynberg, M., Gabrielski, J.Q., Xu, X., Schwarz, B., Zheng, L., Balasubramaniyam, A., Ham, H., Chao, B., et al. (2024). GIMAP5 deficiency reveals a mammalian ceramide-driven longevity assurance pathway. Nature Immunology 25, 282–293. 10.1038/s41590-023-01691-y.

20. Sakaue, S., Kanai, M., Tanigawa, Y., Karjalainen, J., Kurki, M., Koshiba, S., Narita, A., Konuma, T., Yamamoto, K., Akiyama, M., et al. (2021). A cross-population atlas of genetic associations for 220 human phenotypes. Nature Genetics 53, 1415–1424. 10.1038/s41588-021-00931-x.

21. Buniello, A., Suveges, D., Cruz-Castillo, C., Llinares, Manuel B., Cornu, H., Lopez, I., Tsukanov, K., Roldán-Romero, Juan M., Mehta, C., Fumis, L., et al. (2024). Open Targets Platform: facilitating therapeutic hypotheses building in drug discovery. Nucleic Acids Research 53, D1467–D1475. 10.1093/nar/gkae1128.

22. Lee, Y.J., Horie, Y., Wallace, G.R., Choi, Y.S., Park, J.A., Choi, J.Y., Song, R., Kang, Y.M., Kang, S.W., Baek, H.J., et al. (2013). Genome-wide association study identifies GIMAP as a novel susceptibility locus for Behcet’s disease. Ann Rheum Dis 72, 1510–1516. 10.1136/annrheumdis-2011-200288.

23. Groza, T., Gomez, F.L., Mashhadi, H.H., Muñoz-Fuentes, V., Gunes, O., Wilson, R., Cacheiro, P., Frost, A., Keskivali-Bond, P., and Vardal, B. (2023). The International Mouse Phenotyping Consortium: comprehensive knockout phenotyping underpinning the study of human disease. Nucleic acids research 51, D1038–D1045.

24. Gao, E., Lei, Y.H., Shang, X., Huang, Z.M., Zuo, L., Boucher, M., Fan, Q., Chuprun, J.K., Ma, X.L., and Koch, W.J. (2010). A novel and efficient model of coronary artery ligation and myocardial infarction in the mouse. Circulation research 107, 1445–1453.

25. Heineke, J., and Molkentin, J.D. (2006). Regulation of cardiac hypertrophy by intracellular signalling pathways. Nature Reviews Molecular Cell Biology 7, 589–600. 10.1038/nrm1983.

26. Lavine, K.J., Epelman, S., Uchida, K., Weber, K.J., Nichols, C.G., Schilling, J.D., Ornitz, D.M., Randolph, G.J., and Mann, D.L. (2014). Distinct macrophage lineages contribute to disparate patterns of cardiac recovery and remodeling in the neonatal and adult heart. Proceedings of the National Academy of Sciences 111, 16029–16034.

27. Thygesen, K., Alpert, J.S., Jaffe, A.S., Chaitman, B.R., Bax, J.J., Morrow, D.A., White, H.D., and Infarction, E.G.o.b.o.t.J.E.S.o.C.A.C.o.C.A.H.A.W.H.F.T.F.f.t.U.D.o.M. (2018). Fourth universal definition of myocardial infarction (2018). Circulation 138, e618–e651.

28. Abdulla, S., Aevermann, B., Assis, P., Badajoz, S., Bell, S.M., Bezzi, E., Cakir, B., Chaffer, J., Chambers, S., Cherry, J.M., et al. (2025). CZ CELLxGENE Discover: a single-cell data platform for scalable exploration, analysis and modeling of aggregated data. Nucleic Acids Res 53, D886–d900. 10.1093/nar/gkae1142.

29. Malek, A.M., Alper, S.L., and Izumo, S. (1999). Hemodynamic shear stress and its role in atherosclerosis. Jama 282, 2035–2042.

30. Zhang, S.H., Reddick, R.L., Piedrahita, J.A., and Maeda, N. (1992). Spontaneous hypercholesterolemia and arterial lesions in mice lacking apolipoprotein E. Science 258, 468–471.

31. Zarins, C.K., Giddens, D.P., Bharadvaj, B., Sottiurai, V.S., Mabon, R.F., and Glagov, S. (1983). Carotid bifurcation atherosclerosis. Quantitative correlation of plaque localization with flow velocity profiles and wall shear stress. Circulation research 53, 502–514.

32. Ishibashi, S., Goldstein, J.L., Brown, M.S., Herz, J., and Burns, D.K. (1994). Massive xanthomatosis and atherosclerosis in cholesterol-fed low density lipoprotein receptor-negative mice. J Clin Invest 93, 1885–1893. 10.1172/jci117179.

33. Von Scheidt, M., Zhao, Y., Kurt, Z., Pan, C., Zeng, L., Yang, X., Schunkert, H., and Lusis, A.J. (2017). Applications and limitations of mouse models for understanding human atherosclerosis. Cell metabolism 25, 248–261.

34. Nakashima, Y., Plump, A.S., Raines, E.W., Breslow, J.L., and Ross, R. (1994). ApoE-deficient mice develop lesions of all phases of atherosclerosis throughout the arterial tree. Arteriosclerosis and thrombosis: a journal of vascular biology 14, 133–140.

35. Bentzon, J.F., Otsuka, F., Virmani, R., and Falk, E. (2014). Mechanisms of plaque formation and rupture. Circulation research 114, 1852–1866.

36. Pascall, J.C., Webb, L.M., Eskelinen, E.-L., Innocentin, S., Attaf-Bouabdallah, N., and Butcher, G.W. (2018). GIMAP6 is required for T cell maintenance and efficient autophagy in mice. PLoS One 13, e0196504.

37. Muse, E.D., Kramer, E.R., Wang, H., Barrett, P., Parviz, F., Novotny, M.A., Lasken, R.S., Jatkoe, T.A., Oliveira, G., Peng, H., et al. (2017). A Whole Blood Molecular Signature for Acute Myocardial Infarction. Sci Rep 7, 12268. 10.1038/s41598-017-12166-0.

38. Guo, S., Wu, J., Zhou, W., Liu, X., Liu, Y., Zhang, J., Jia, S., Li, J., and Wang, H. (2021). Identification and analysis of key genes associated with acute myocardial infarction by integrated bioinformatics methods. Medicine 100, e25553.

39. Li, X., He, X., Zhang, Y., Hao, X., Xiong, A., Huang, J., Jiang, B., Tong, Z., Huang, H., Yi, L., and Chen, W. (2025). Uncovering Hippo pathway-related biomarkers in acute myocardial infarction via scRNA-seq binding transcriptomics. Scientific Reports 15, 10368. 10.1038/s41598-025-94820-6.

40. Hu, P., Liu, J., Zhao, J., Wilkins, B.J., Lupino, K., Wu, H., and Pei, L. (2018). Single-nucleus transcriptomic survey of cell diversity and functional maturation in postnatal mammalian hearts. Genes & development 32, 1344–1357.

41. Kalluri, A.S., Vellarikkal, S.K., Edelman, E.R., Nguyen, L., Subramanian, A., Ellinor, P.T., Regev, A., Kathiresan, S., and Gupta, R.M. (2019). Single-cell analysis of the normal mouse aorta reveals functionally distinct endothelial cell populations. Circulation 140, 147–163.

42. Rahkonen, O., Su, M., Hakovirta, H., Koskivirta, I., Hormuzdi, S.G., Vuorio, E., Bornstein, P., and Penttinen, R. (2004). Mice With a Deletion in the First Intron of the Col1a1 Gene Develop Age-Dependent Aortic Dissection and Rupture. Circulation Research 94, 83–90. 10.1161/01.RES.0000108263.74520.15.

43. Seto, T., Yamamoto, T., Shimojima, K., and Shintaku, H. (2017). A novel COL1A1 mutation in a family with osteogenesis imperfecta associated with phenotypic variabilities. Human Genome Variation 4, 1–3.

44. Hamczyk, M.R., Nevado, R.M., Gonzalo, P., Andrés-Manzano, M.J., Nogales, P., Quesada, V., Rosado, A., Torroja, C., Sánchez-Cabo, F., Dopazo, A., et al. (2024). Endothelial-to-Mesenchymal Transition Contributes to Accelerated Atherosclerosis in Hutchinson-Gilford Progeria Syndrome. Circulation 150, 1612–1630. 10.1161/CIRCULATIONAHA.123.065768.

45. Hahn, C., and Schwartz, M.A. (2009). Mechanotransduction in vascular physiology and atherogenesis. Nature reviews Molecular cell biology 10, 53–62.

46. Kuhlencordt, P.J., Gyurko, R., Han, F., Scherrer-Crosbie, M., Aretz, T.H., Hajjar, R., Picard, M.H., and Huang, P.L. (2001). Accelerated Atherosclerosis, Aortic Aneurysm Formation, and Ischemic Heart Disease in Apolipoprotein E/Endothelial Nitric Oxide Synthase Double-Knockout Mice. Circulation 104, 448–454. 10.1161/hc2901.091399.

47. Kunjathoor, V.V., Febbraio, M., Podrez, E.A., Moore, K.J., Andersson, L., Koehn, S., Rhee, J.S., Silverstein, R., Hoff, H.F., and Freeman, M.W. (2002). Scavenger receptors class AI/II and CD36 are the principal receptors responsible for the uptake of modified low density lipoprotein leading to lipid loading in macrophages. Journal of Biological Chemistry 277, 49982–49988.

48. Cury, J., Haudiquet, M., Trejo, V.H., Mordret, E., Hanouna, A., Rotival, M., Tesson, F., Bonhomme, D., Ofir, G., and Quintana-Murci, L. (2024). Conservation of antiviral systems across domains of life reveals immune genes in humans. Cell Host & Microbe 32, 1594-1607. e1595.

49. Chawla, A., Barak, Y., Nagy, L., Liao, D., Tontonoz, P., and Evans, R.M. (2001). PPAR-γ dependent and independent effects on macrophage-gene expression in lipid metabolism and inflammation. Nature medicine 7, 48–52.

50. Rahaman, S.O., Lennon, D.J., Febbraio, M., Podrez, E.A., Hazen, S.L., and Silverstein, R.L. (2006). A CD36-dependent signaling cascade is necessary for macrophage foam cell formation. Cell metabolism 4, 211–221.

51. Mocci, G., Sukhavasi, K., Örd, T., Bankier, S., Singha, P., Arasu, U.T., Agbabiaje, O.O., Mäkinen, P., Ma, L., and Hodonsky, C.J. (2024). Single-cell gene-regulatory networks of advanced symptomatic atherosclerosis. Circulation research 134, 1405–1423.

52. Rogacev, K.S., Cremers, B., Zawada, A.M., Seiler, S., Binder, N., Ege, P., Große-Dunker, G., Heisel, I., Hornof, F., and Jeken, J. (2012). CD14++ CD16+ monocytes independently predict cardiovascular events: a cohort study of 951 patients referred for elective coronary angiography. Journal of the American College of Cardiology 60, 1512–1520.

53. Kuda, O., Pietka, T.A., Demianova, Z., Kudova, E., Cvacka, J., Kopecky, J., and Abumrad, N.A. (2013). Sulfo-N-succinimidyl oleate (SSO) inhibits fatty acid uptake and signaling for intracellular calcium via binding CD36 lysine 164: SSO also inhibits oxidized low density lipoprotein uptake by macrophages. Journal of Biological Chemistry 288, 15547–15555.

54. Else, P.L. (2020). The highly unnatural fatty acid profile of cells in culture. Progress in Lipid Research 77, 101017. 10.1016/j.plipres.2019.101017.

55. Spector, A.A., Mathur, S.N., and Kaduce, T.L. (1980). Lipid nutrition and metabolism of cultured mammalian cells. Progress in Lipid Research 19, 155–186. 10.1016/0163-7827(80)90003-X.

56. The, G.C., Aguet, F., Anand, S., Ardlie, K.G., Gabriel, S., Getz, G.A., Graubert, A., Hadley, K., Handsaker, R.E., Huang, K.H., et al. (2020). The GTEx Consortium atlas of genetic regulatory effects across human tissues. Science 369, 1318–1330. 10.1126/science.aaz1776.

57. Heng, T.S.P., Painter, M.W., Elpek, K., Lukacs-Kornek, V., Mauermann, N., Turley, S.J., Koller, D., Kim, F.S., Wagers, A.J., Asinovski, N., et al. (2008). The Immunological Genome Project: networks of gene expression in immune cells. Nature Immunology 9, 1091–1094. 10.1038/ni1008-1091.

58. Shadur, B., Asherie, N., Kfir-Erenfeld, S., Dubnikov, T., NaserEddin, A., Schejter, Y.D., Elpeleg, O., Mor-Shaked, H., and Stepensky, P. (2021). A human case of GIMAP6 deficiency: a novel primary immune deficiency. European Journal of Human Genetics 29, 657–662.

59. Jurgens, S.J., Wang, X., Choi, S.H., Weng, L.-C., Koyama, S., Pirruccello, J.P., Nguyen, T., Smadbeck, P., Jang, D., and Chaffin, M. (2024). Rare coding variant analysis for human diseases across biobanks and ancestries. Nature Genetics 56, 1811–1820.

60. Nikpay, M., Goel, A., Won, H.-H., Hall, L.M., Willenborg, C., Kanoni, S., Saleheen, D., Kyriakou, T., Nelson, C.P., Hopewell, J.C., et al. (2015). A comprehensive 1000 Genomes–based genome-wide association meta-analysis of coronary artery disease. Nature Genetics 47, 1121–1130. 10.1038/ng.3396.

61. Biobank, M.B.A.L.W.S.J.W.V.A.M.J.G.C.M.S., and 15, A.o.U.R.D.P.T.C.S.H.h.o.o.-.-.-W.X.h.o.o.-.-.-R.E.A. (2024). Genomic data in the all of us research program. Nature 627, 340–346.

62. Damrauer, S.M., Chaudhary, K., Cho, J.H., Liang, L.W., Argulian, E., Chan, L., Dobbyn, A., Guerraty, M.A., Judy, R., Kay, J., et al. (2019). Association of the V122I Hereditary Transthyretin Amyloidosis Genetic Variant With Heart Failure Among Individuals of African or Hispanic/Latino Ancestry. JAMA 322, 2191–2202. 10.1001/jama.2019.17935.

63. McKay, G.J., Silvestri, G., Chakravarthy, U., Dasari, S., Fritsche, L.G., Weber, B.H., Keilhauer, C.N., Klein, M.L., Francis, P.J., Klaver, C.C., et al. (2011). Variations in apolipoprotein E frequency with age in a pooled analysis of a large group of older people. Am J Epidemiol 173, 1357–1364. 10.1093/aje/kwr015.

64. Verma, A., Huffman, J.E., Rodriguez, A., Conery, M., Liu, M., Ho, Y.-L., Kim, Y., Heise, D.A., Guare, L., and Panickan, V.A. (2024). Diversity and scale: Genetic architecture of 2068 traits in the VA Million Veteran Program. Science 385, eadj1182.

65. Wang, Q., Dhindsa, R.S., Carss, K., Harper, A.R., Nag, A., Tachmazidou, I., Vitsios, D., Deevi, S.V.V., Mackay, A., Muthas, D., et al. (2021). Rare variant contribution to human disease in 281,104 UK Biobank exomes. Nature 597, 527–532. 10.1038/s41586-021-03855-y.

66. Marjamaa, J., Tulamo, R., Abo-Ramadan, U., Hakovirta, H., Frösen, J., Rahkonen, O., Niemelä, M., Bornstein, P., Penttinen, R., and Kangasniemi, M. (2006). Mice with a deletion in the first intron of the Col1a1 gene develop dissection and rupture of aorta in the absence of aneurysms: High-resolution magnetic resonance imaging, at 4.7 T, of the aorta and cerebral arteries. Magnetic Resonance in Medicine: An Official Journal of the International Society for Magnetic Resonance in Medicine 55, 592–597.

67. Tian, K., Xu, Y., Sahebkar, A., and Xu, S. (2020). CD36 in atherosclerosis: pathophysiological mechanisms and therapeutic implications. Current atherosclerosis reports 22, 59.

68. Glatz, J.F., Heather, L.C., and Luiken, J.J. (2024). CD36 as a gatekeeper of myocardial lipid metabolism and therapeutic target for metabolic disease. Physiological reviews 104, 727–764.

## REFERENCES

3. Libby, P. (2021). The changing landscape of atherosclerosis. Nature 592, 524–533.

8. Limoges, M.-A., Cloutier, M., Nandi, M., Ilangumaran, S., and Ramanathan, S. (2021). The GIMAP Family Proteins: An Incomplete Puzzle. Frontiers in Immunology 12. 10.3389/fimmu.2021.679739.

9. Yao, Y., Du Jiang, P., Chao, B.N., Cagdas, D., Kubo, S., Balasubramaniyam, A., Zhang, Y., Shadur, B., NaserEddin, A., and Folio, L.R. (2022). GIMAP6 regulates autophagy, immune competence, and inflammation in mice and humans. Journal of Experimental Medicine 219.

10. Park, A.Y., Leney-Greene, M., Lynberg, M., Gabrielski, J.Q., Xu, X., Schwarz, B., Zheng, L., Balasubramaniyam, A., Ham, H., Chao, B., et al. (2024). GIMAP5 deficiency reveals a mammalian ceramide-driven longevity assurance pathway. Nature Immunology 25, 282–293. 10.1038/s41590-023-01691-y.

11. Sakaue, S., Kanai, M., Tanigawa, Y., Karjalainen, J., Kurki, M., Koshiba, S., Narita, A., Konuma, T., Yamamoto, K., Akiyama, M., et al. (2021). A cross-population atlas of genetic associations for 220 human phenotypes. Nature Genetics 53, 1415–1424. 10.1038/s41588-021-00931-x.

12. Buniello, A., Suveges, D., Cruz-Castillo, C., Llinares, Manuel B., Cornu, H., Lopez, I., Tsukanov, K., Roldán-Romero Juan M., Mehta, C., Fumis, L., et al. (2024). Open Targets Platform: facilitating therapeutic hypotheses building in drug discovery. Nucleic Acids Research 53, D1467–D1475. 10.1093/nar/gkae1128.

13. Lee, Y.J., Horie, Y., Wallace, G.R., Choi, Y.S., Park, J.A., Choi, J.Y., Song, R., Kang, Y.M., Kang, S.W., Baek, H.J., et al. (2013). Genome-wide association study identifies GIMAP as a novel susceptibility locus for Behcet’s disease. Ann Rheum Dis 72, 1510–1516. 10.1136/annrheumdis-2011-200288.

14. Gao, E., Lei, Y.H., Shang, X., Huang, Z.M., Zuo, L., Boucher, M., Fan, Q., Chuprun, J.K., Ma, X.L., and Koch, W.J. (2010). A novel and efficient model of coronary artery ligation and myocardial infarction in the mouse. Circulation research 107, 1445–1453.

15. Lavine, K.J., Epelman, S., Uchida, K., Weber, K.J., Nichols, C.G., Schilling, J.D., Ornitz, D.M., Randolph, G.J., and Mann, D.L. (2014). Distinct macrophage lineages contribute to disparate patterns of cardiac recovery and remodeling in the neonatal and adult heart. Proceedings of the National Academy of Sciences 111, 16029–16034.

16. Malek, A.M., Alper, S.L., and Izumo, S. (1999). Hemodynamic shear stress and its role in atherosclerosis. Jama 282, 2035–2042.

17. Zhang, S.H., Reddick, R.L., Piedrahita, J.A., and Maeda, N. (1992). Spontaneous hypercholesterolemia and arterial lesions in mice lacking apolipoprotein E. Science 258, 468– 471.

18. Zarins, C.K., Giddens, D.P., Bharadvaj, B., Sottiurai, V.S., Mabon, R.F., and Glagov, S. (1983). Carotid bifurcation atherosclerosis. Quantitative correlation of plaque localization with flow velocity profiles and wall shear stress. Circulation research 53, 502–514.

19. Emini Veseli, B., Perrotta, P., De Meyer, G.R.A., Roth, L., Van der Donckt, C., Martinet, W., and De Meyer, G.R.Y. (2017). Animal models of atherosclerosis. European Journal of Pharmacology 816, 3–13. 10.1016/j.ejphar.2017.05.010.

20. Nakashima, Y., Plump, A.S., Raines, E.W., Breslow, J.L., and Ross, R. (1994). ApoE-deficient mice develop lesions of all phases of atherosclerosis throughout the arterial tree. Arteriosclerosis and thrombosis: a journal of vascular biology 14, 133–140.

21. Bentzon, J.F., Otsuka, F., Virmani, R., and Falk, E. (2014). Mechanisms of plaque formation and rupture. Circulation research 114, 1852–1866.

22. Hu, P., Liu, J., Zhao, J., Wilkins, B.J., Lupino, K., Wu, H., and Pei, L. (2018). Single-nucleus transcriptomic survey of cell diversity and functional maturation in postnatal mammalian hearts. Genes & development 32, 1344–1357.

23. Kalluri, A.S., Vellarikkal, S.K., Edelman, E.R., Nguyen, L., Subramanian, A., Ellinor, P.T., Regev, A., Kathiresan, S., and Gupta, R.M. (2019). Single-cell analysis of the normal mouse aorta reveals functionally distinct endothelial cell populations. Circulation 140, 147–163.

24. Kunjathoor, V.V., Febbraio, M., Podrez, E.A., Moore, K.J., Andersson, L., Koehn, S., Rhee, J.S., Silverstein, R., Hoff, H.F., and Freeman, M.W. (2002). Scavenger receptors class AI/II and CD36 are the principal receptors responsible for the uptake of modified low density lipoprotein leading to lipid loading in macrophages. Journal of Biological Chemistry 277, 49982–49988.

25. Shadur, B., Asherie, N., Kfir-Erenfeld, S., Dubnikov, T., NaserEddin, A., Schejter, Y.D., Elpeleg, O., Mor-Shaked, H., and Stepensky, P. (2021). A human case of GIMAP6 deficiency: a novel primary immune deficiency. European Journal of Human Genetics 29, 657–662.

26. Tarhan, L., Bistline, J., Chang, J., Galloway, B., Hanna, E., and Weitz, E. (2023). Single Cell Portal: an interactive home for single-cell genomics data. BioRxiv.

27. Chen, P.-Y., Qin, L., and Simons, M. (2022). Imaging and analysis of oil red o-stained whole aorta lesions in an aneurysm hyperlipidemia mouse model. Journal of visualized experiments: JoVE, 10.3791/61277.

28. Pascall, J.C., Webb, L.M., Eskelinen, E.-L., Innocentin, S., Attaf-Bouabdallah, N., and Butcher, G.W. (2018). GIMAP6 is required for T cell maintenance and efficient autophagy in mice. PLoS One 13, e0196504.

29. Toda, G., Yamauchi, T., Kadowaki, T., and Ueki, K. (2021). Preparation and culture of bone marrow-derived macrophages from mice for functional analysis. STAR protocols 2, 100246.

30. Pirruccello, J.P., Chaffin, M.D., Chou, E.L., Fleming, S.J., Lin, H., Nekoui, M., Khurshid, S., Friedman, S.F., Bick, A.G., Arduini, A., et al. (2022). Deep learning enables genetic analysis of the human thoracic aorta. Nature Genetics 54, 40–51. 10.1038/s41588-021-00962-4.

